# Protein sequence sampling and prediction from structural data

**DOI:** 10.1101/2021.09.06.459171

**Authors:** Gabriel A. Orellana, Javier Caceres-Delpiano, Roberto Ibañez, Michael P. Dunne, Leonardo Alvarez

## Abstract

The increasing integration between protein engineering and machine learning has led to many interesting results. A problem still to solve is to evaluate the likelihood that a sequence will fold into a target structure. This problem can be also viewed as sequence prediction from a known structure.

In the current work, we propose improvements in the recent architecture of Geometric Vector Perceptrons [1] in order to optimize the sampling of sequences from a known backbone structure. The proposed model differs from the original in that there is: (i) no updating in the vectorial embedding, only in the scalar one, (ii) only one layer of decoding. The first aspect improves the accuracy of the model and reduces the use of memory, the second allows for training of the model with several tasks without incurring data leakage.

We treat the trained classifier as an Energy-Based Model and sample sequences by sampling amino acids in a non-autoreggresive manner in the empty positions of the sequence using energy-guided criteria and followed by random mutation optimization. We improve the median identity of samples from 40.2% to 44.7%.

An additional question worth investigating is whether sampled and original sequences fold into similar structures independent of their identity. We chose proteins in our test set whose sampled sequences show low identity (under 30%) but for which our model predicted favorable energies. We used AlphaFold [2, 3] and observed that the predicted structures for sampled sequences highly resemble the predicted structures for original sequences, with an average TM-score of 0.84.

## 1 Introduction

Proteins are one of the most interesting and highly studied macromolecules due to their diverse functional features. Protein sequences strictly determine the spontaneous folding that characterize their 3D structures. Different groups have shown applications of Deep Learning (DL) methods to major topics in Biochemistry. An example of this is the protein folding problem, making it possible to predict structure of proteins just from amino acid sequences [4, 3]. A relatively new approach for modeling proteins involves the use of graphs. Proteins can be seen as points in the space connected by different kinds of interactions. This information can be represented in a graph that consists of a set of nodes and a set of edges composed by pairs of nodes. [5] proposed an architecture based on Graph Convolutions, leading to new architectures named Graph Convolutional Networks (GCN).

The goal of the current work is to develop a GCN model and adjacent algorithms that address the problem of amino acid sequence prediction and sequence sampling given the structure of a backbone. This work is inspired by existing work [6, 1], but prioritizing the optimization of sampled sequences.

Similar works in the literature [6, 1] have trained their models in the task of Autoregressive Single Amino acid Prediction (ARSAP), consisting of predicting the current amino acid on a protein sequence, given the real protein structure and the real sequence prior to the current amino acid. ARSAP and identity between original and sampled sequences are the commonly used metrics. Previously used sample procedure [6] consists of predicting amino acids in an autoregressive (AR) manner (similar to ARSAP but with cumulative error). We propose to consider the logits in our model as a negative energy in the same fashion as it is used in Energy-Based Models (EBM) [7], and then iteratively sample the amino acid with the lowest energy until all the sequence is filled.

Finally, we use AlphaFold [2, 3] to predict the structure of some of our sampled sequences and show that even in cases when identities are low, if energy of sequences is low, the structure of the sampled sequence predicted by AlphaFold is highly similar to the predicted structure of the original sequence.

The contributions of this work are four: *(i)* we propose a new GCN layer architecture taking as starting point the one proposed by [6] along with the improvements made by [1], increasing the performance in ARSAP and sequence sampling; *(ii)* using the advantages given by the proposed architecture, we train our model with different auxiliary tasks, improving its performance in the metric of sampling identity; *(iii)* by exploiting some characteristics of our architecture we implement an energy-guided sampling procedure that outperformed the AR procedure in the sequence identity metric; *(iv)* we show that our model is able to generate sequences that fold into structures highly similar to the target structures even in cases when these sequences are not similar.

## 2 Methods

### Graph features

Following several works in modeling proteins as graphs [6, 8, 9, 10] we considered the use of node features, accounting for the features of single amino acids, and edge features, accounting for interactions between them. Based on the importance of close interactions in proteins, [6] proposes that each node keeps the connections to its *k*-nearest neighbors in atomic distance. We performed tests to evaluate the optimal value of this meta-parameter and found it was *k* = 35.

Based on the recent success in the use of 3D vectors as complementary features for the this problem, presented in [1], we include two extra sets of features that corresponds to the 3D vector representation of *(i)* the normalized distances between all the atoms in the backbone for each amino acid, included as node features; and *(ii)* the normalized distances between the *Cα* of each amino acid and the *Cα* of the *k* nearest neighbors, included as edge features.

The four kinds of features (vectorial and scalar node features plus vectorial and scalar edge features) and the amino acid sequence are then encoded using a standard trainable embedding and introduced into the model, which is trained to output one value per each possible amino acid (20 in total) for each position in the sequence following a classification paradigm.

### Architecture

Inspired by the graph network application to this problem shown in [6] and the improvements made in [1], we propose a GCN with four layers of only structural information (encoding layers) and a single layer of both structural and sequence information (decoding layer). This design decision differs from the architectures previously cited, where the number of encoding and decoding layers were the same. We proposed this change expecting: *(i)* more dependence on structural information. We expect the performance to decrease on the ARSAP task, but increase or maintain in the sequence sampling task, since the latter relies heavily on structural information while the former uses both structure and sequence information; *(ii)* more flexibility in the training of our model, since we have just one decoding layer, there is no problem of data leakage with masks different that AR mask; and *(iii)* more speed in sampling because, for each step in the process the inputs of the decoding layer are the same and, therefore, just that layer needs to be executed.

### Graph layer

The functioning of layers used in our architecture is described in pseudocode in appendix A and displayed in figure 1,

**Figure 1:**
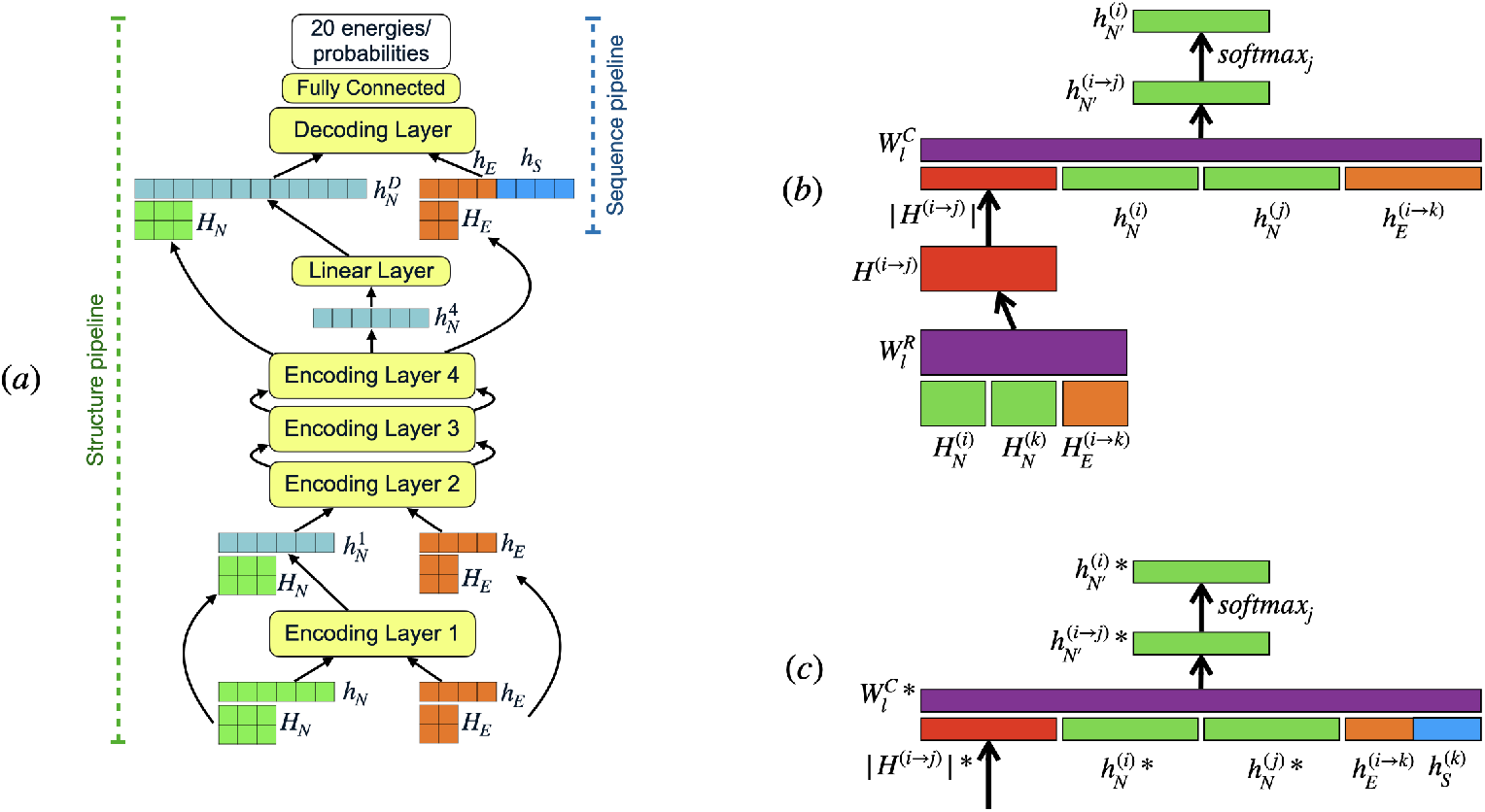
**(a)** Full network diagram integrating Encoder and Decoder layers from feature inputs to probability/energy **(b)** General architecture of an encoder layer, and **(c)** General architecture of decoder layer. Node features on green, edge features on orange, vector convolution output on red, sequence embeddings on blue, trainable weights on purple. Parameters in decoder layer with a * have a dimension of double the size as in encoder layers. In (a), length of pipelines for structure and sequence information can be seen on the sides of the full network diagram.

### Loss functions

Under the regular probabilistic setting, the model is trained to approximate the probabilistic distributions of the target. Considering the use of the softmax function, equation 1 describes the loss function *L*_1_, with *q*_*a*_ as the logits for class *a*. In case of the EBM, the loss function attempts to minimize the energy of the target class while increasing the energies of the rest. Here we use a function *L*_2_ from the family of Contrastive Free Energy losses [11] (equation 2).

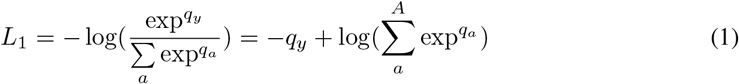

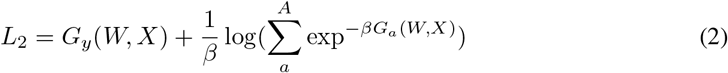

Equations 1 and 2 are very similar, differing in the sign on the energy factors. We therefore use the logits trained in the standard probabilistic setting as negative energy in an EBM setting.

### Tasks

We describe the main task used in previous works [6, 1] as well as the 3 auxiliary tasks employed in this work in appendix B.

### Evaluation of Energy for Sequence-Structure Tuple (E_SST_)

We define a formula for calculating the energy of a sequence and a structure using our model (see appendix C and figure C.1).

### Sampling procedure

We believe that the AR sampling algorithm used in literature [6, 1] does not take advantage of two things: *(i)* As we described previously, normal sampling is inefficient because it needs to run the entire forward of the protein at every step. In order to deal with this we modified the architecture to have just one decoding layer and modified the sampling procedure to run the encoding part of the forward just once. *(ii)* AR sampling forces to sample the next position in the sequence instead of other positions that may have less energy (more certainty). Based on this, we develop a non-AR Energy-Based sample procedure (see appendix D). It is worth noting that we do not see big differences in performance regarding identity between AR and non-AR sampling (table S3). In the discussion section we explore the utility of optimizing the energy.

## 3 Results

Following the setting of [6] and also adopted [1], i.e. ARSAP task with metaparameter *k* = 30, we evaluate the performance of the three models in table 1. Regarding the performance of our model with the proposed additions, table S2 displays the results of our model trained in the main task and the addition of the auxiliary tasks described in the previous section, with a parameter *k* = 35. The model trained with all the 4 tasks proposed achieves a perplexity of 5.34 and an accuracy of 47.2%. On the other hand, table S4 shows the results on the Sequence Sampling task, comparing the model trained with the different tasks and using AR and non-AR sampling procedures. The non-AR procedure using the model trained with 4 tasks achieves a median identity on the whole test set of 44.3% and just on short sequences of 35.1%. For calculating metrics in tables S2 and S4, we removed proteins with His-tag from training, validation and test data sets (around 20%). The effect of this modification can be seen in table S1. For comparison sake, our model obtained a median identity of 44.7% when trained and tested on the original data set (with the His-tag proteins).

**Table 1:** Results in AR Single Amino acid Prediction (ARSAP) and Sequence Sampling with metaparameter k=30 and trained with main task only. Sampling results are shown in median value (more details in table S3). ^1^: Model proposed in [6]; ^2^: Model proposed in [1]

|  | ARSAP |  | Sampling |  |
| --- | --- | --- | --- | --- |
|  | Perplexity | Accuracy (%) | Identity short seqs (%) | Identity total (%) |
| AttGraph<sup>1</sup> | 6.93 | 39.4 | 28.3 | 36.4 |
| GVP<sup>2</sup> | 5.29 | 46.6 | 32.2 | 40.3 |
| Ours | 5.43 | 46.6 | 34.6 | 43.9 |

## 4 Discussion

Based on our interpretation of the logits as the negative of the energy, a natural point of interest is whether the energy of sampled sequences is correlated with identity. We found a high correlation between these two variables with a Pearson’s *r* of −0.8778 (figure S1).

An important question unaddressed in previous works is whether sampled sequences could exhibit low levels of identity with the original sequence and still generate 3D foldings with high similarity with the input structure. There is evidence of some structural similarity between foldings of sequences with identities of 20% [12]. If these cases occur, it would imply that the model is able to learn how to produce a similar structure using different patterns of amino acids. Following the correlation shown in figure S1, we tested whether the energy of sequences can be used as a predictor of structural similarity. In order to obtain an anecdotal insight of this phenomena, we selected a subset of 6 proteins in the test set where their sampled sequences have the lowest energy and their sequence identities are lower than 30%. Later, we used AlphaFold [2, 3] to predict the structure of the sampled sequences and compare them with the original ones (see figure 2 and table S5, details in appendix E). We obtained an average TM-score of 0.84 ± 0.20 with a median of 0.92 comparing the original and sampled structures. As a control, we followed the same procedure with other 6 proteins that matched the same criterion for identity, but with higher energies (table S6). The best candidate sequences for proteins on this subset obtained an average TM-score of 0.64 ± 0.16 with a median of 0.72. From this we can conclude that the energy obtained from our model shows encouraging potential as predictor of performance of the model as well as likelihood function between pairs of sequences and structures (see figure S2), although further work is still needed, such as structure predictions for bigger sets of sampled sequences or alternative experimental settings.

**Figure 2:**
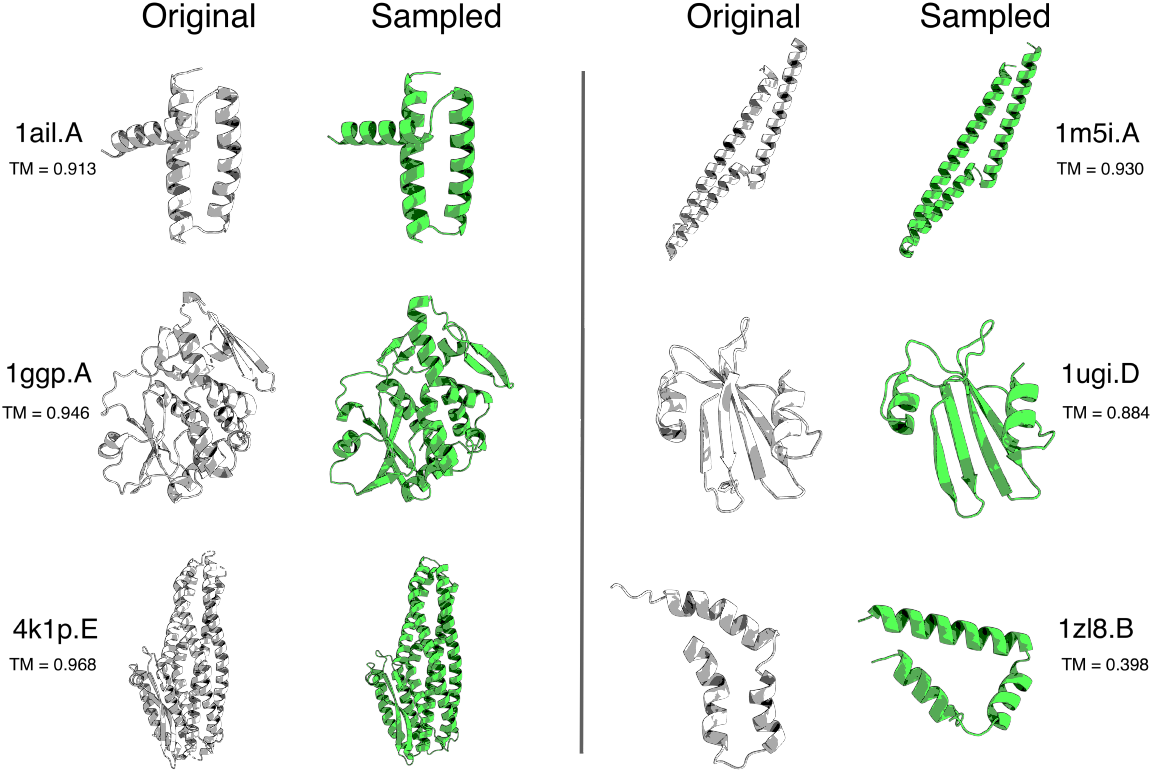
Alignment of structures generated by AlphaFold from original (white) and sampled (green) sequences. PDB codes and chains are shown for each structure, as well as the calculated TM-score between original and sampled structures.

## Appendices

### A Graph convolutional layer

The functioning of layers used in our architecture is described on pseudocode in algorithm A.1 and displayed in figure 1.b and 1.c, where *h*_*N*_ are the scalar node features and *h*_*E*_ are scalar edge features, both ∈ ℝ^*S*^, while *H*_*N*_ and *H*_*E*_ are the vectorial node and edge features respectively, both ∈ ℝ^*V* ×3^. being *S* the hidden dimension of scalar features and *V* the hidden dimension of vectorial features.

**Algorithm A.1:**
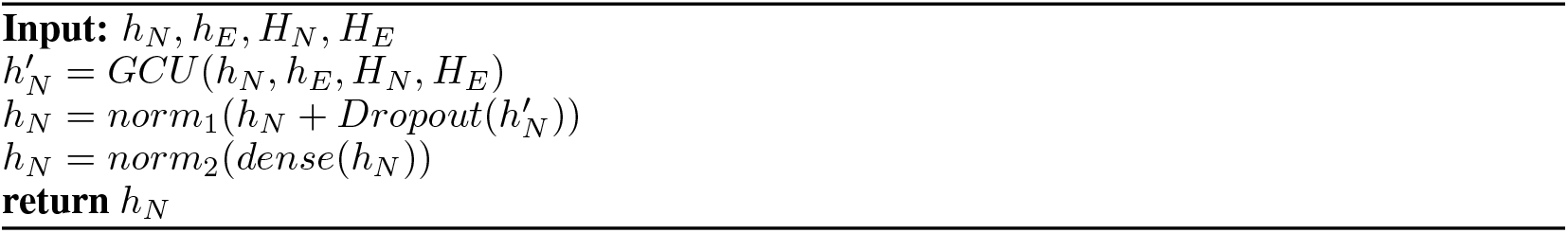
Graph convolutional layer

With *GCU* being the Graph convolutional unit, *norm*_1_ and *norm*_2_ are standard normalization layers with trainable terms for gain and bias; and *dense* is a standard Position-wise Feedforward layer used as a dense layer. As it can be seen, in contrast with [1], our model does not update both scalar and vectorial node features in each layer, but just the scalar node features. This also allows us to keep the standard dense layers used in [6]. The GCU is shown in pseudocode on algorithm A.2, where *K* is the hyperparameter for the number of nearest neighbors nodes considered in the elaboration of the graph, while *W*_*R*_ ∈ ℝ^*S*×(3*V*)^, *W*_*C*_ ∈ ℝ^(4*S*)×*S′*^ and *W*_*out*_ ∈ ℝ^*S′*×*S*^ are weight matrices with *S*′ as an intermediate hidden layer dimension.

**Algorithm A.2:**
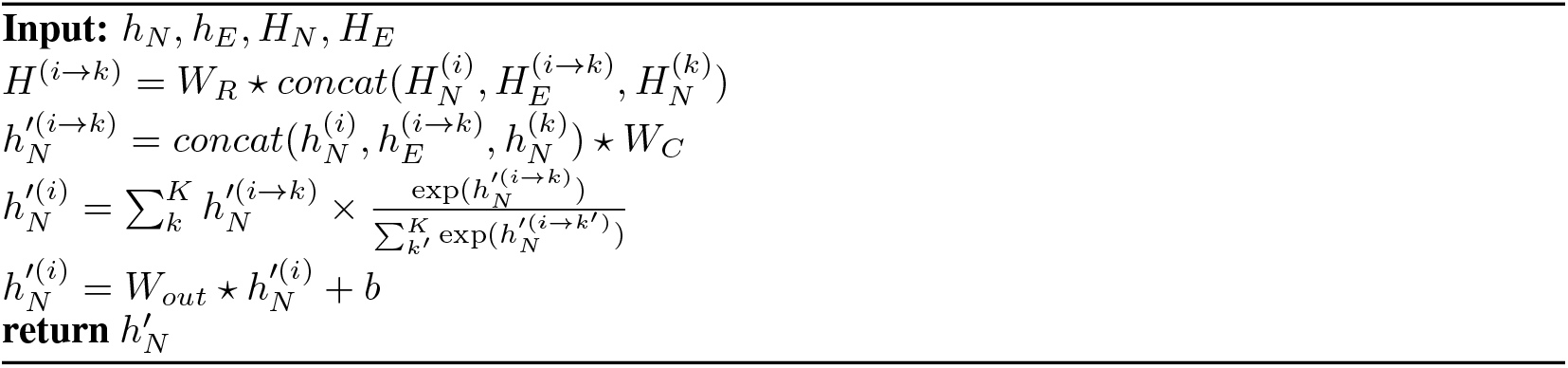
Graph convolutional unit

For the decode layer, the sequence information 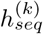 is concatenated to 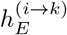 and then introduced into the regular data flow (see figure 1.b). The hidden dimension of scalar features is then duplicated in order to fit the size of the data input using a standard linear layer.

### B Tasks

The regular task associated with this problem and already described in [6] is the autoregressive single amino acid prediction, that can be stated as:

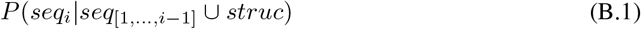

Where *seq*_*i*_ is the sequence in the *i* position. In notation of EBM, we can describe it as:

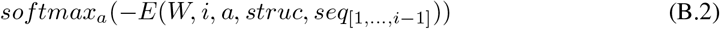

In our study we employ three additional tasks:

1. Knowing the following amino acids in the sequence, but no the previous ones (inverse autoregressive prediction): *P*(*seq*_*i*_|*seq*_[*i*+1,...,*N*]_ ∪ *struc*) or *softmax*_*a*_(−*E*(*W, i, a, struc, seq*_[*i*+1*,...,N*]_))
2. Knowing all the amino acids in the sequence except the current one: *P*(*seq*_*i*_|*seq* − {*seq*_*i*_} ∪ *struc*) or *softmax*_*a*_(−*E*(*W, i, a, struc, seq* − {*seq*_*i*_}))
3. Knowing none of the amino acids in the sequence: *P*(*seq*_*i*_|*struc*) or *softmax*_*a*_(−*E*(*W, i, a, struc,* ∅))

**Figure C.1:**
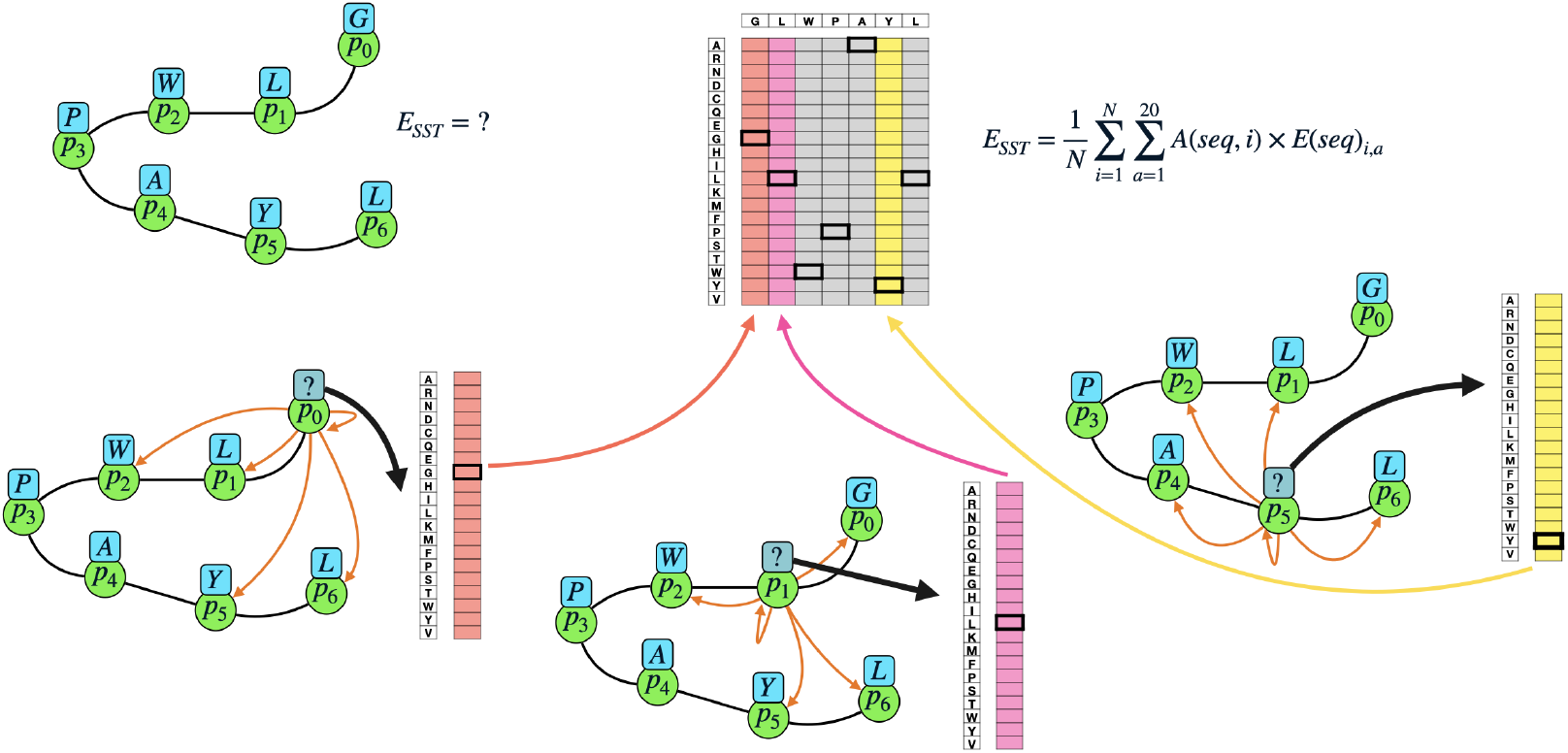
Illustration of E_SST_. 20 energies are calculated for each spot of the sequence using as inputs of the model the structure and the sequence with a mask on the amino acid in said spot. Then, from those 20 energies, the energy of the amino acid in said spot is considered to calculate the average.

It is worth mentioning that the three additional tasks can be easily employed in the training of our model due to the decision of having just one decoding layer (see figure 1.b). In particular, auxiliary task number 2 would create data leakage in the training process in case that the decoding architecture would consist in more than one layer since the whole batch containing the possible edge connection of one or more proteins is executed concurrently.

### C E_SST_

We consider the tuple of a structure *struc* and a sequence *seq*. We run the tuple through the model in evaluation mode in the same configuration as in the auxiliary task 2. We define function for belonging to sequence *seq* in position *i* for an amino acid *a* as

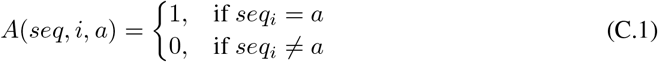

Then we define:

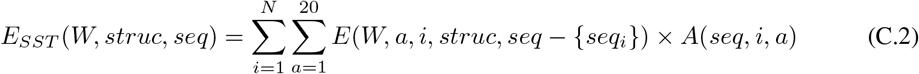

Being the summation of energies only from amino acids in sequence *seq* given its immediate surrounding in sequence and structure, as it is illustrated in figure C.1.

It is worth mentioning that this procedure is the same as the auxiliary task 2, therefore, as it was explained in section B, it needs to have a single decoding layer.

### D Energy-Based Sampling

The initial step in the Energy-Based Sampling procedure is to start from an empty sequence and sample single amino acids based on their energy in a non-AR manner. We call this procedure **Energy-Guided Sampling** (EGS) and it is explained in algorithm D.1.

**Algorithm D.1:**
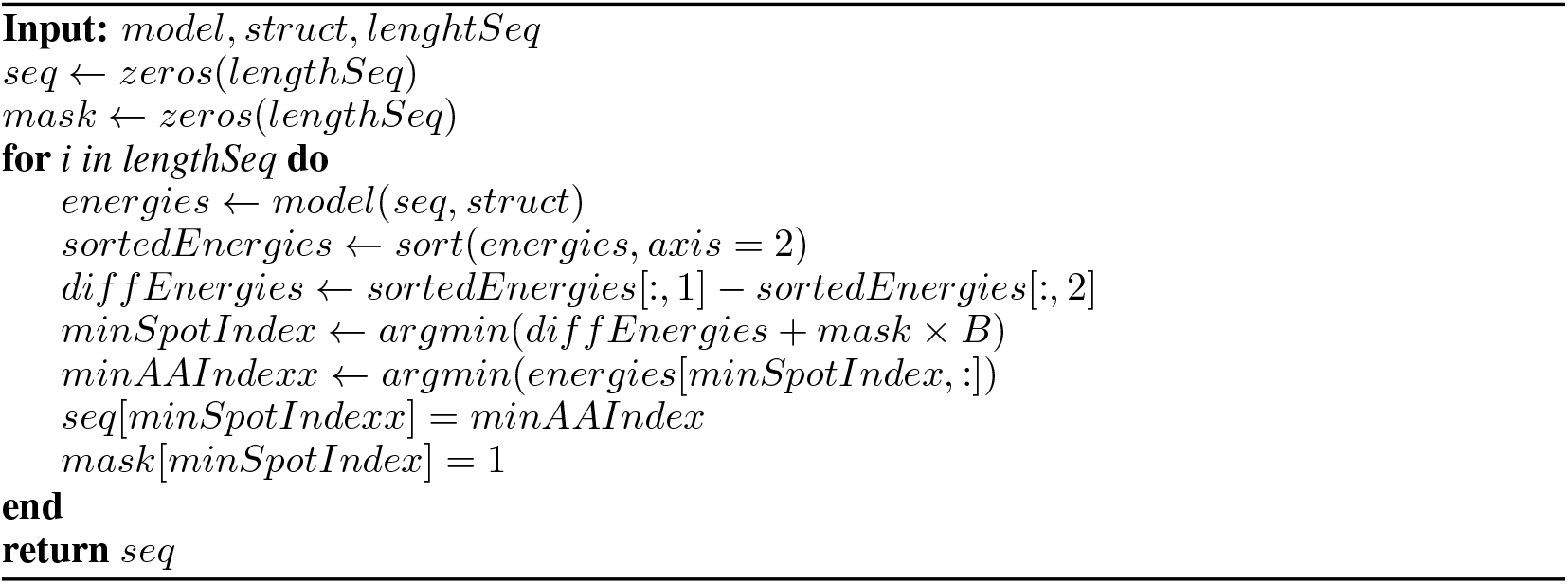
Energy-Guided Sampling

Following EGS we expect that, according to the model, the sampled amino acids are optimal given the surroundings at the point when they are sampled. On the downside, once all the amino acids are sampled, it is not guaranteed that amino acids are optimal given their surroundings since they changed from the moment where the amino acids were sampled. Considering this we employ a step of **Energy-Guided Mutation Optimization** (see algorithm D.2).

**Algorithm D.2:**
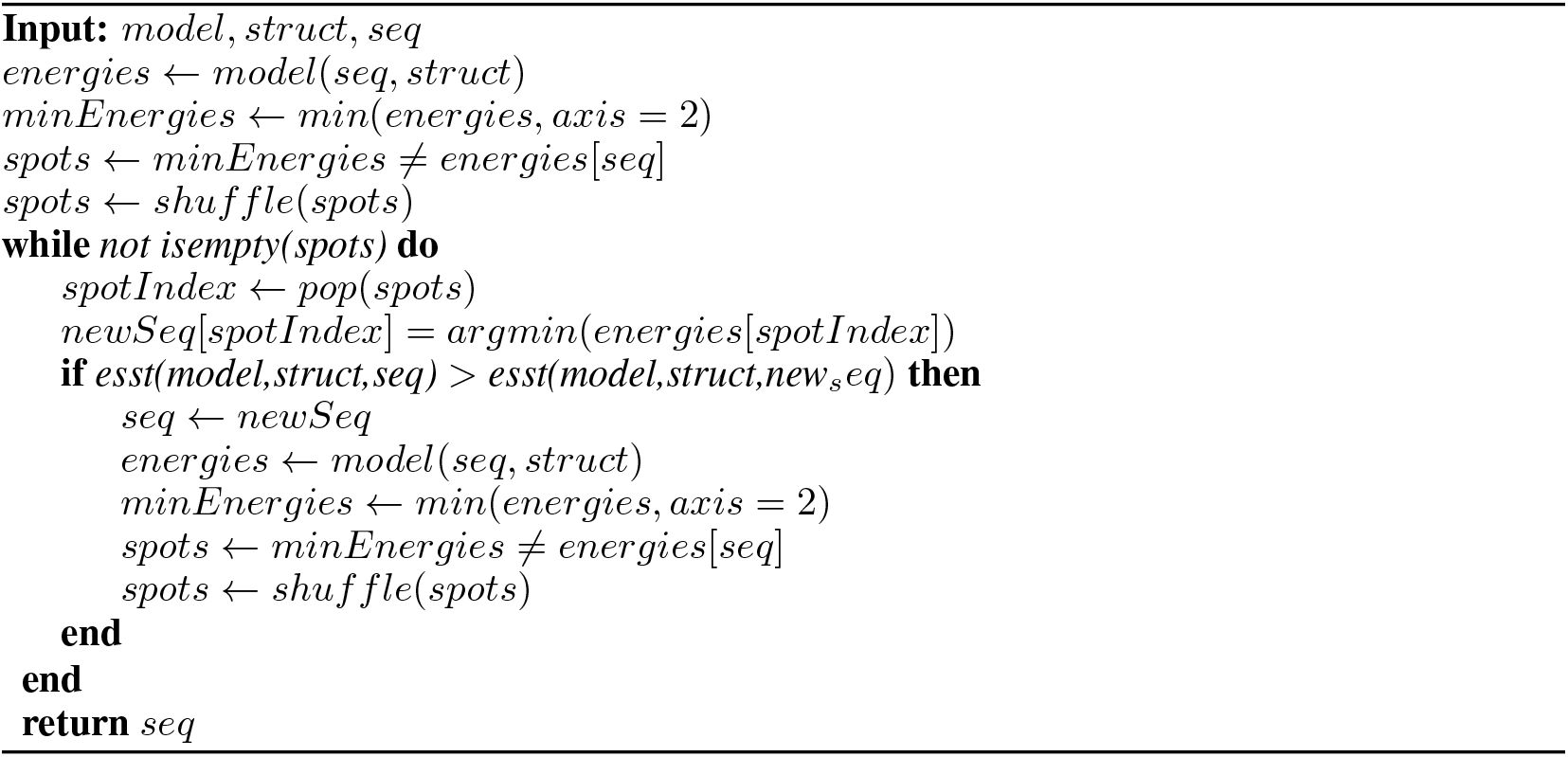
Energy-Guided Mutation Optimization

After the two steps of Energy-Guided Sampling and Optimization, the optimality of the energy of sequence is still not guaranteed. We implement a final step of Random Mutation Optimization (RMO), when we generate a high number (*N* = 1000) of random single mutations and evaluate whether they improve the energy of the sequence employing E_SST_.

### E Parameters for AlphaFold predictions

MMSeqs2 was run with *s* = 7.5, databases were BFD (consensus only), MGnify, Uniclust30 (2018_08), UniRef90. Duplicate sequences were discarded. The resulting MSA was used to search the PDB70 database for templates. AlphaFold was run on this MSA and these templates using all available models as of August 2021, and the highest ranked PDB according to pLDDT was selected.

## Supplementary information

**Figure S1:**
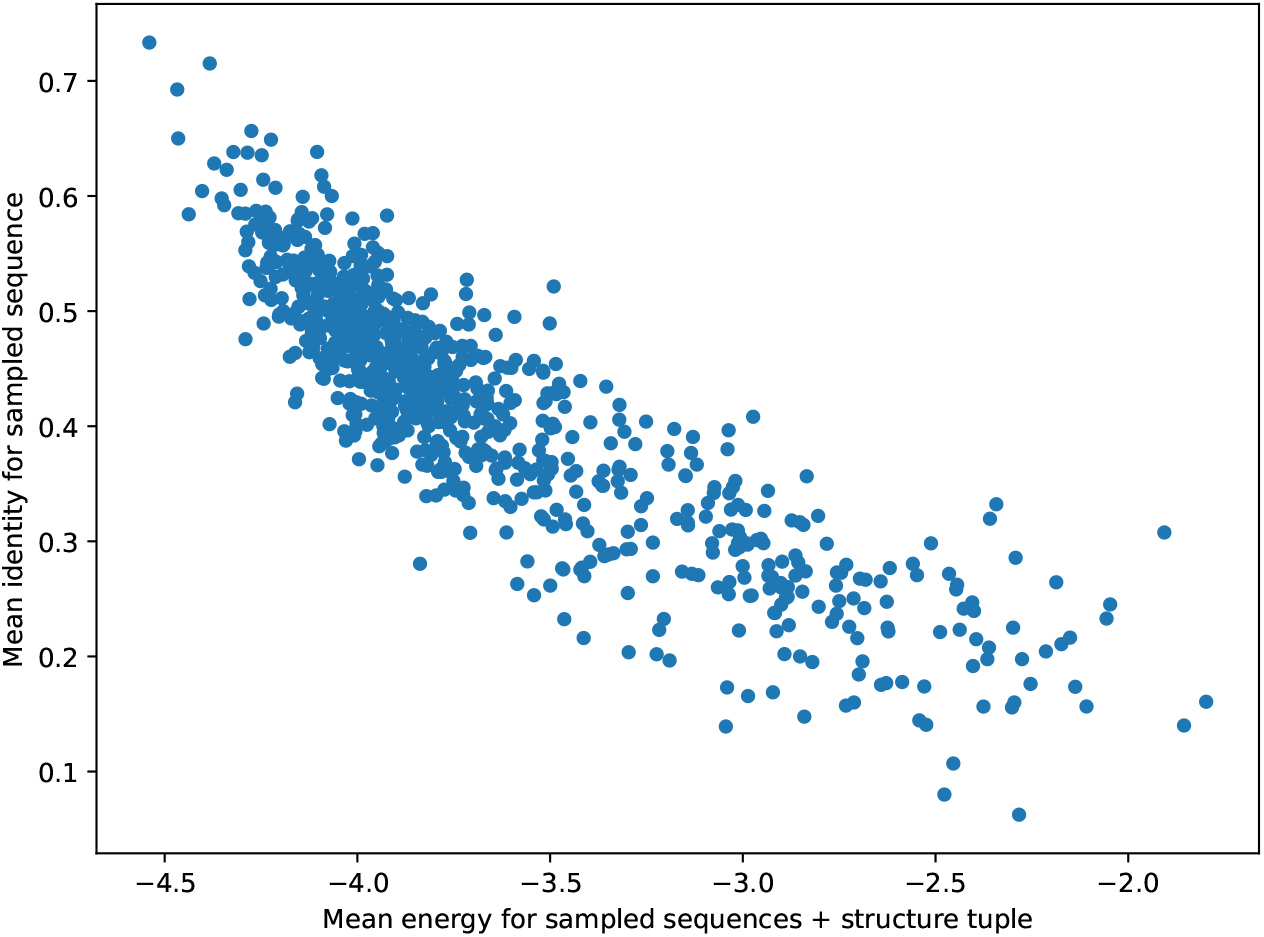
Mean protein identity all sampled sequences for every protein in the test data set compared to their Energy of Sequence-Structure Tuple (E_SST_). Pearson’s r of −0.8778.

**Table S1:** Results on both tasks comparing final version of our model trained and tested in the CATH dataset [13] with and without proteins with His-tag. Model were trained with main (ARSAP) task only. Accuracy and identities are presented in percentage.

|  | ARSAP |  | Sampling |  |
| --- | --- | --- | --- | --- |
|  | Perplexity | Accuracy | Ident. short seqs | Ident. total |
| Ours with His-tag proteins | 5.40 | 47.0 | 34.0 | 41.7 |
| Ours without His-tag proteins | 5.52 | 46.4 | 33.4 | 41.2 |

**Table S2:** Results in Autoregressive Single Amino Acid Prediction. Our model was trained in the main task and an increasing number of auxiliary tasks. Meta parameter *k* = 35. Databases for this measure were modified, removing all the protein with His-tag from training, validation and test subsets.

| Training | Perplexity | Accuracy (%) |
| --- | --- | --- |
| Main task only | 5.52 | 46.4 |
| Main + aux task 1 | 5.64 | 46.2 |
| Main + aux tasks 1, 2 | 5.40 | 46.7 |
| Main + aux tasks 1, 2, 3 | 5.44 | 46.6 |

**Table S3:**
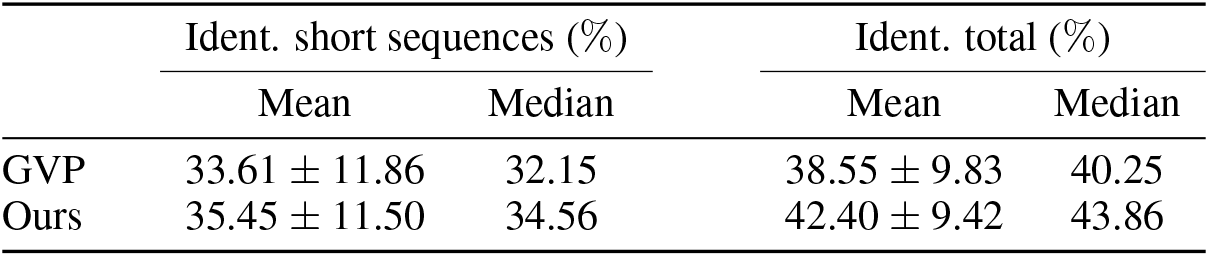
Results in Autoregressive (AR) or non-Autoregressive (nAR) sequence sampling tasks. Model was trained in the main task and an increasing number of auxiliary tasks. Meta parameter *k* = 35. Databases for this measure were modified, removing all the protein with His-tag from training, validation and test subsets.

|  | Ident. short sequences (%) |  | Ident. total (%) |  |
| --- | --- | --- | --- | --- |
|  | Mean | Median | Mean | Median |
| GVP | 33.61 ± 11.86 | 32.15 | 38.55 ± 9.83 | 40.25 |
| Ours | 35.45 ± 11.50 | 34.56 | 42.40 ± 9.42 | 43.86 |

**Table S4:** Results in Autoregressive (AR) or non-Autoregressive (nAR) sequence sampling tasks. Model was trained in the main task and an increasing number of auxiliary tasks. Meta parameter *k* = 35. Databases for this measure were modified, removing all the protein with His-tag from training, validation and test subsets.

|  | Ident. total (%) |  | Energy |  |
| --- | --- | --- | --- | --- |
|  | Mean | Median | Mean | Median |
| AR main task only | 41.5 ± 9.9 | 43.1 | -3.61 ± 0.49 | -3.76 |
| nAR main task only | 41.2 ± 10.1 | 42.5 | -3.79 ± 0.49 | -3.97 |
| AR main + aux task 1 | 41.8 ± 9.8 | 43.5 | -3.69 ± 0.49 | -3.85 |
| nAR main + aux task 1 | 41.6 ± 10.1 | 43.2 | -3.84 ± 0.50 | -4.01 |
| AR main + aux tasks 1, 2 | 42.1 ± 9.6 | 43.7 | -3.43 ± 0.52 | -3.59 |
| nAR main + aux tasks 1, 2 | 42.1 ± 10.0 | 43.9 | -3.56 ± 0.53 | -3.74 |
| AR main + aux tasks 1, 2, 3 | 42.3 ± 10.1 | 43.8 | -3.50 ± 0.52 | -3.68 |
| nAR main + aux tasks 1, 2, 3 | 42.3 ± 10.6 | 44.1 | -3.69 ± 0.50 | -3.86 |

**Table S5:** Statistics of structures predicted with AlphaFold from sampled sequences sampled selected for exhibiting low energy among with low identity. Statistics correspond to the best candidate sequence according to the lowest energy criterion for all the sampled sequences for that protein.

| Protein | Length | Energy | Identity (%) | pLDDT | TM-Score |
| --- | --- | --- | --- | --- | --- |
| 1ugi.D | 82 | -3.758 | 26.83 | 90.83 | 0.884 |
| 2guz.B | 65 | -3.582 | 29.63 | 70.49 | 0.398 |
| 1ggp.A | 234 | -3.543 | 27.95 | 92.28 | 0.968 |
| 3zbh.A | 90 | -3.536 | 29.52 | 90.48 | 0.930 |
| 1q90.B | 212 | -3.502 | 27.14 | 90.14 | 0.914 |
| 1ce7.A | 241 | -3.482 | 26.07 | 86.32 | 0.946 |
| Average | | | | | 0.840 ± 0.200 |
| Median |  |  |  |  | 0.922 |

**Table S6:** Statistics of structures predicted with AlphaFold from sampled sequences sampled selected for exhibiting medium or high energy among with low identity. Statistics correspond to the best candidate sequence according to the lowest energy criterion for all the sampled sequences for that protein. The energy criterion for selecting these proteins was 6 proteins in the upper half of the energy distribution and with the same distance (in the distribution) between each other.

| Protein | Length | Energy | Identity (%) | pLDDT | TM-Score |
| --- | --- | --- | --- | --- | --- |
| 1b35.D | 57 | -2.970 | 29.82 | 75.15 | 0.674 |
| 3cra.A | 239 | -2.838 | 27.20 | 60.34 | 0.805 |
| 3j7y.d | 162 | -2.757 | 24.69 | 75.38 | 0.889 |
| 1ifw.A | 92 | -2.638 | 27.17 | 87.24 | 0.558 |
| 1urf.A | 81 | -2.482 | 22.22 | 87.83 | 0.760 |
| 2l35.A | 62 | -1.971 | 25.81 | 72.16 | 0.416 |
| Average | | | | | $0.684 \pm 0.158$ |
| Median |  |  |  |  | 0.717 |

**Figure S2:**
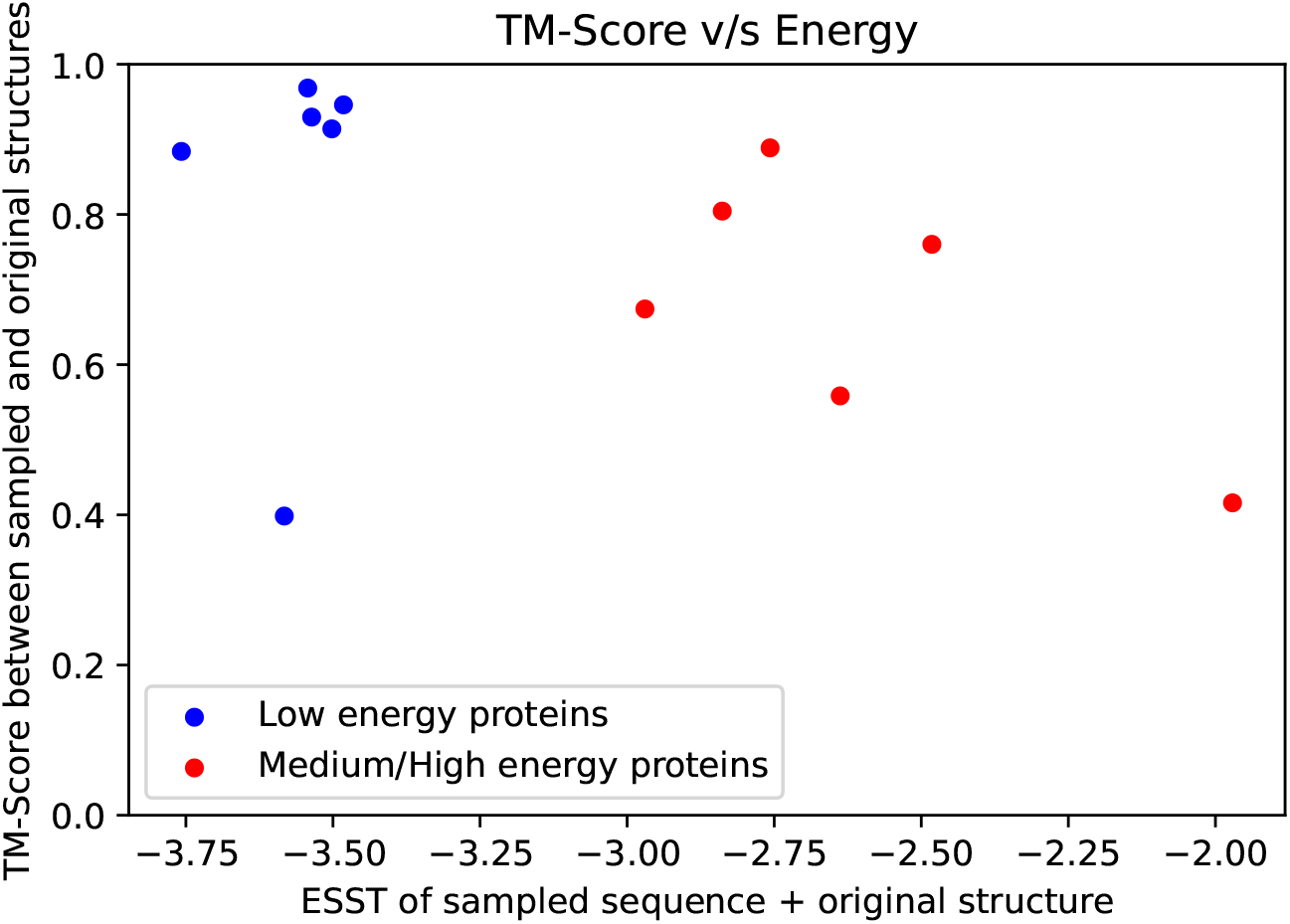
Energy of sampled sequence and original structure v/s TM-Score between sampled structure (predicted with AlphaFold) and original structure, comparing subsets with low energies from table S5 (in blue) and medium/high energies from table S6 (in red).

## Notes

### Competing Interest Statement

Provisional patent applications have been filed based on the results presented here.

### Summary of Updates

Michael P. Dunne was added as author. We added an illustration of how the energy of a sequence is calculated. We added stats on the performance of the two sampling schemes (AR and non-AR). Selection of proteins to be evaluated by structure prediction was changed, choosing the lowest energies on an individual sequence basis instead of average of sequences per protein.

